# Analysis of global and site-specific radiation damage in cryo-EM

**DOI:** 10.1101/258178

**Authors:** Johan Hattne, Dan Shi, Calina Glynn, Chih-Te Zee, Marcus Gallagher-Jones, Michael W. Martynowycz, Jose A. Rodriguez, Tamir Gonen

**Author notes:** To whom correspondence should be addressed: T.G.

## Abstract

Micro-crystal electron diffraction (MicroED) is an emerging method in cryo-EM for structure determination using nanocrystals. It has been used to solve structures of a diverse set of biomolecules and materials, in some cases to sub-atomic resolution. However, little is known about the damaging effects of the electron beam on samples during such measurements. We assess global and site-specific damage from electron radiation on nanocrystals of proteinase K and of a prion hepta-peptide and find that the dynamics of electron-induced damage follow well-established trends observed in X-ray crystallography. Metal ions are perturbed, disulfide bonds are broken, and acidic side chains are decarboxylated while the diffracted intensities decay exponentially with increasing exposure. A better understanding of radiation damage in MicroED improves our assessment and processing of all types of cryo-EM data.

## Introduction

Structure determination relies on interpreting the outcome of interactions of a beam of quanta with a sample. Most quanta pass through samples without any interaction at all (Henderson, 1995). Other quanta scatter either elastically, whereby they interact with the sample without losing any energy, or inelastically by depositing part of their energy to the sample. In a conventional diffraction measurement, the information that can be gained increases with the number of elastically scattered quanta. Inelastic scattering events manifest as damage, introduce noise, and ultimately limit the signal that can be extracted from the sample.

The success of crystallographic structure determination ultimately depends on the ratio of elastic to inelastic scattering events (Nave & Hill, 2005). While this ratio is greater in electron diffraction than in X-ray diffraction (Henderson, 1995), a single incident electron carries sufficient energy to knock out several electrons from an atom in an inelastic scattering event. These ejected, secondary electrons and their associated Auger electrons are mobile even at 77 K (Jones et al., 1987), and can, depending on the chemical composition of the sample and its surrounding mother liquor, cause additional ionization and excitation events in the crystal (Garman, 2010). Additional damage from thermal diffusion of atomic and molecular radicals produced by these ionization events is curbed by keeping the sample at cryogenic temperatures, where diffusion is limited (Henderson & Unwin, 1975; Hayward & Glaeser, 1979; Uyeda et al., 1980; Jeng & Chiu, 1984). The absorbed energy, which is related to radiation damage, depends on the chemical composition of the crystal and the medium in which it is embedded, the temperature at which the measurement is performed, and the energy of the incident electrons.

As damage accumulates, its effects become apparent at specific sites as well as throughout the entire crystal. As crystalline order deteriorates, the fraction of unit cells contributing to crystalline diffraction decreases, the *B*-factor generally increases (Kmetko et al., 2006), and the observed unit cell volume may increase as the lattice expands (Ravelli et al., 2002). Consequently, the signal of the obtained diffraction pattern, which varies with the square of the number of scattering unit cells, decreases. To some extent, damage can be compensated for by appropriate scaling procedures (Diederichs, 2006), and may be further mitigated in nanocrystals as the probability that secondary electrons escape before causing further damage is higher than in large crystals (Nave & Hill, 2005; Sanishvili et al., 2011).

Site-specific damage may be observed if the impact of radiation damage on the crystal is not uniformly distributed but more selective towards certain moieties (Weik et al., 2000). Unlike global damage, site-specific damage can only be assessed once the dataset has been integrated and phased, and a real-space density map is calculated. Site-specific damage becomes apparent when certain bonds are more susceptible to damage than others, and may remain invisible if it only occurs in a small fraction of the unit cells or if it is masked by phases calculated from an undamaged model.

Exposure of the sample to the electron beam results in immediate damage even at cryogenic temperatures as in electron cryo-electron microscopy (cryo-EM). Low-dose procedures helped minimize the exposure to the sample prior to data recording (Uyeda et al., 1980). Early studies indicated that organic samples deteriorate 4–5× faster at room temperature compared with cryogenic temperatures (Unwin & Henderson, 1975; Hayward & Glaeser, 1979; Jeng & Chiu, 1984). In these studies, either two-dimensional crystals and/or three-dimensional crystals were used in electron diffraction to look at the overall decay of reflections following exposure to the electron beam. These studies indicated that atomic resolution information (defined at better than 3 Å) was lost after exposure of the sample to only 3 e^−^ Å^−2^. In 2013, a new method for cryo-EM was unveiled and termed MicroED, or 3D electron crystallography of microscopic crystals (Shi et al., 2013; Nannenga et al., 2014b). With continuous rotation MicroED, which is the preferred method of data collection in MicroED, significant loss of diffraction intensity was observed at resolutions better than ~3 Å when only ~3 e^−^ Å^−2^ were used, consistent with past studies (Hayward & Glaeser, 1979; Jeng & Chiu, 1984; Baker et al., 2010).

Owing to the strong interaction of electrons with matter, and the fact that only diffraction data are collected (no imaging), high-resolution structures can be determined by MicroED from three-dimensional nanocrystals with significantly less total electron exposure than what is normally used for other cryo-EM modalities. Recent MicroED experiments demonstrated that complete data sets could be collected from a single nanocrystal using a total exposure of less than ~1–2 e^−^ Å^−2^ (Nannenga et al., 2014a; de la Cruz et al., 2017), making it possible to design new and improved experiments to test for beam induced damage to the specimen with increasing exposure to the electron beam.

In this study, we set out to determine the damaging effects of electron radiation using MicroED and nanocrystals of a well characterized sample, proteinase K, and a short hepta-peptide with a bound metal. MicroED data were collected using exposures of 0.0017–0.007 e^−^ Å^−2^ s^−1^. Such low exposure allowed us to repeatedly measure the same wedge of reciprocal space from the same crystal and to compare the data obtained over increasing exposure as the experiment progressed. The data was also sufficiently complete to allow us to investigate the effects not only in reciprocal space but also in real space, such that both global and site-specific damage can be observed. The data indicate that beam damage is a limiting factor in high-resolution cryo-EM methods, and the results have implications for all EM methods that use considerably higher electron exposures for imaging.

## Results

### Indicators of global damage

For both the globular proteinase K and hepta-peptide samples, the overall weakening of the diffraction spots resulting from the loss of crystalline order can be modeled by exponential decay as a function of absorbed dose (Blake & Phillips, 1962; Liebschner et al., 2015) (Figure 1). We note that this model appears to systematically underestimate the intensities of the weakest reflections at high exposures for all samples. This observation does not necessarily invalidate the model: weak reflections are difficult to measure accurately due to noise; to the extent these reflections can be measured at all; they are not discernible by eye. Profiles derived from stronger surrounding reflections are likely to overestimate their intensities, and outlier rejection due to, for example, ill-fitting background models introducing further bias toward more intense reflections.

**Figure 1:**
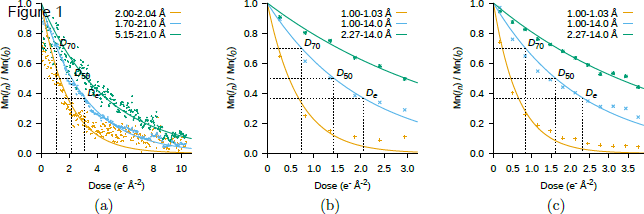
Exposure dependency of the mean intensity of the unscaled integrated reflections. (a) proteinase K, (b) the hepta-peptide, GSNQNNF, recorded at a exposure rate of 0.0028 e^−^ Å^−2^ s^−1^, and (c) GSNQNNF at 0.0017 e^−^ Å^−2^ s^−1^. Spots at high resolution fade significantly faster than spots at low resolution. For proteinase K, the electron exposure at which the mean intensity across the entire recorded resolution range is reduced to 70%, 50%, and *e*^−1^ of its extrapolated value at zero dose is estimated to be *D*_70_ = 1.1 e^−^ Å^−2^, *D*_50_ = 2.2 e^−^ Å^−2^, and *D_e_* = 3.1 e^−^ Å^−2^. The corresponding exposures for the 0.0028 e^−^ Å^−2^ s^−1^ and 0.0017 e^−^ Å^−2^ s^−1^ peptide datasets are *D*_70_ = 0.73 e^−^ Å^−2^, *D*_50_ = 1.4 e^−^ Å^−2^, *D_e_* = 2.1 e^−^ Å^−2^ and *D*_70_ = 0.83 e^−^ Å^−2^, *D*_50_ = 1.6 e^−^ Å^−2^, *D_e_* = 2.3 e^−^ Å^−2^, respectively.

After an exposure of 1 e^−^ Å^−2^ the average intensity of all observed reflections in proteinase K has decreased to 73% of its extrapolated value at zero dose (Figure 1). After an additional 1.6 e^−^ Å^−2^, the high-resolution limit of the data has dropped from 1.7 Å to 1.9 Å (Table 1). Similar trends can be seen in the peptide images at high and low exposure rates where the intensities have dropped to 61% and 65%, respectively, by 1 e^−^ Å^−2^. However, the highest-resolution reflections in the peptide data are much stronger than those in proteinase K and the effect of exposure on optical resolution is consequently not as pronounced. For both peptide data sets, the optical resolution decreases to ~1.1 Å by 2 e-Å^−2^ (Table 2, Table 3).

**Table 1.**
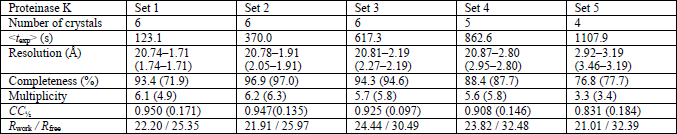
Processing and refinement statistics for proteinase K. The exposure time of a multi-crystal dataset is denoted <*t*_exp_>; it is defined as the mean cumulative irradiated time of all the frames in the dataset. Numbers in parentheses refer to the highest-resolution shell for either merging or refinement. Owing to varying response to radiation, damage-induced non-isomorphism causes the number of datasets that can be merged to decrease at the higher exposures.

**Table 2.** Processing and refinement statistics for the hepta-peptide GSNQNNF at 0.0028 e^−^ Å^−2^ s^−1^. Rows as per Table 1.

| GSNQNNF<br>($0.0028 \text{ e}^- \text{ Å}^{-2} \text{ s}^{-1}$) | Set 1 | Set 2 | Set 3 | Set 4 | Set 5 | Set 6 |
| --- | --- | --- | --- | --- | --- | --- |
| Number of crystals | 10 | 10 | 10 | 10 | 3 | 3 |
| $\langle t_{\text{exp}} \rangle$ (s) | 96.7 | 289.1 | 480.1 | 670.1 | 860.5 | 1050.5 |
| Resolution (Å) | 13.95–1.01<br>(1.03–1.01) | 13.92–1.01<br>(1.03–1.01) | 13.94–1.01<br>(1.03–1.01) | 14.01–1.15<br>(1.19–1.15) | 13.85–1.26<br>(1.48–1.26) | 6.53–1.37<br>(1.48–1.37) |
| Completeness (%) | 80.7 (72.5) | 81.6 (71.9) | 82.4 (68.9) | 83.4 (86.1) | 56.1 (56.6) | 63.5 (61.1) |
| Multiplicity | 6.9 (6.2) | 6.9 (6.5) | 6.6 (5.4) | 6.5 (6.6) | 3.3 (3.3) | 2.7 (2.8) |
| $CC_{1/2}$ | 98.8 (91.1) | 98.9 (91.5) | 98.6 (60.0) | 98.9 (79.1) | 96.1 (44.6) | 3.3 (14.2) |
| $R_{\text{work}} / R_{\text{free}}$ | 17.49 / 15.18 | 18.88 / 18.18 | 24.68 / 22.82 | 26.36 / 29.87 | 36.54 / 32.75 | 36.45 / 43.44 |

**Table 3.** Processing and refinement statistics for the hepta-peptide GSNQNNF at 0.0017 e^−^ Å ^−2^ s^−1^. Rows as per Table 1.

| GSNQNNF<br>($0.0017 \text{ e}^- \text{ Å}^{-2} \text{ s}^{-1}$) | Set 1 | Set 2 | Set 3 | Set 4 | Set 5 | Set 6 | Set 7 | Set 8 | Set 9 | Set 10 | Set 11 | Set 12 |
| --- | --- | --- | --- | --- | --- | --- | --- | --- | --- | --- | --- | --- |
| Number of crystals | 12 | 12 | 12 | 12 | 5 | 12 | 11 | 12 | 12 | 11 | 11 | 11 |
| $\langle t_{\text{exp}} \rangle$ (s) | 97.2 | 291.3 | 484.6 | 677.1 | 869.0 | 1060.6 | 1251.0 | 1442.9 | 1636.2 | 1832.2 | 2024.3 | 2214.0 |
| Resolution (Å) | 11.48–1.01<br>(1.03–1.01) | 13.97–1.01<br>(1.03–1.01) | 13.89–1.01<br>(1.03–1.01) | 13.89–1.02<br>(1.03–1.02) | 13.86–1.01<br>(1.03–1.01) | 13.86–1.01<br>(1.03–1.01) | 13.70–1.13<br>(1.15–1.13) | 13.76–1.16<br>(1.20–1.16) | 13.71–1.21<br>(1.26–1.21) | 13.72–1.31<br>(1.38–1.31) | 13.67–1.46<br>(1.59–1.46) | 13.68–1.45<br>(1.57–1.45) |
| Completeness (%) | 79.4 (72.7) | 79.2 (71.4) | 79.0 (70.8) | 76.3 (64.4) | 74.3 (67.0) | 77.8 (64.0) | 74.4 (76.8) | 78.6 (81.2) | 80.8 (81.0) | 79.4 (79.9) | 77.5 (78.5) | 79.8 (67.6) |
| Multiplicity | 8.6 (7.5) | 8.4 (6.5) | 8.5 (7.0) | 8.5 ( 7.2) | 3.7 (3.3) | 8.1 (5.1) | 7.5 (7.7) | 7.8 (8.3) | 7.7 (6.8) | 7.4 ( 6.7) | 7.7 (7.9) | 7.6 (7.9) |
| $CC_{1/2}$ | 98.3 (92.0) | 98.2 (92.8) | 98.0 (83.3) | 98.4 (66.5) | 98.2 (37.3) | 95.7 (44.3) | 86.2 (70.1) | 98.2 (75.0) | 89.5 (69.9) | 95.8 (60.1) | 93.9 (31.9) | 89.9 (71.1) |
| $R_{\text{work}} / R_{\text{free}}$ | 17.92 /<br>16.78 | 18.15 /<br>16.80 | 18.67 /<br>17.69 | 19.82 /<br>20.49 | 23.60 /<br>24.47 | 24.55 /<br>26.72 | 21.49 /<br>22.13 | 24.70 /<br>30.84 | 27.49 /<br>31.61 | 24.14 /<br>33.40 | 23.22 /<br>35.72 | 23.84 /<br>32.74 |

The highest resolution reflections, those providing atomic resolution, are most sensitive to disruption of crystalline order. Consequently, as a result of subtle long-range changes in the lattice, weak, high-resolution reflections fade into the background faster than strong, low-resolution reflections (Blake & Phillips, 1962; Howells et al., 2009), and the highest observable resolution decreases with dose (Table 1, Table 2, Table 3). In conjunction, fine features in real space density maps disappear at a higher rate than the overall molecular envelope. Generally, high resolution information, defined as better than 2 Å, was significantly decayed when exposures greater than ~3 e^−^ Å^−2^ were used. Beyond ~4 e^−^ Å^−2^, the reflections at a resolution finer than 2 Å have dropped to 10% of their extrapolated value at zero dose in diffraction patterns of proteinase K. The corresponding exposure for the peptide at high and low dose rates are similar at 2.7 e^−^ Å^−2^ and 3.2 e^−^ Å^−2^, respectively. As an additional proxy for global damage by electrons we use changes in relative *B*-factor, *B*_rel_ (Kmetko et al., 2006), calculated over the reflections in the resolution range common to all datasets of a given sample and exposure rate. For all but the poorest diffracting crystals, *B*_rel_ increases monotonically with absorbed dose (Figure 2). We also find that the unit cell volume, although a much less reliable indicator of global radiation damage, generally increases with exposure (Murray & Garman, 2002). In our measurements, the unit cell volumes of proteinase K and of the peptide increased by a modest 1.8% from 452 to 460 nm^3^ and 1.15 o 1.17 nm^3^, respectively.

**Figure 2:**
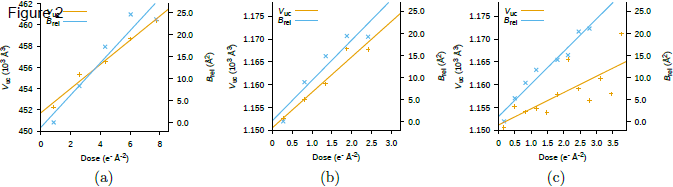
Exposure dependency of the unit cell volume, *V*_uc_, and relative *B*-factor, *B*_rel_. *V*_uc_ and *B*_rel_ were averaged across all the crystals at each exposure. For *B*_rel_ only reflections in a sufficiently large resolution range common to all datasets were considered (20.8– 3.20 Å for proteinase K; 14.0–1.2 Å for GSNQNNF).

### Site-specific damage

Localized chemical changes within the macromolecule can be analyzed by observing changes in the density attributed to specific atoms in real space. We observe site-specific damage even at exposures as small as 0.1 e^−^ Å^−2^. For example, the presence of positive *m*F_o_-*D*F_c_ difference density is detected around the sulfur atoms of the disulfide bonds in proteinase K, indicating that the disulfide bridge was breaking apart in a significant fraction of the unit cells (Helliwell, 1988) even at total exposure <0.9 e^−^ Å^−2^ (Figure 3). At this exposure, the overall diffraction intensity was reduced to 75% of its extrapolated value at zero dose, indicating that 86% of the unit cells are still diffracting to high resolution (Blake & Phillips, 1962). As the exposure increases, the positive difference density is replaced by negative difference density in the location of the bond and the 2*m*F_o_-*D*F_c_ density progressively weakens.

**Figure 3:**
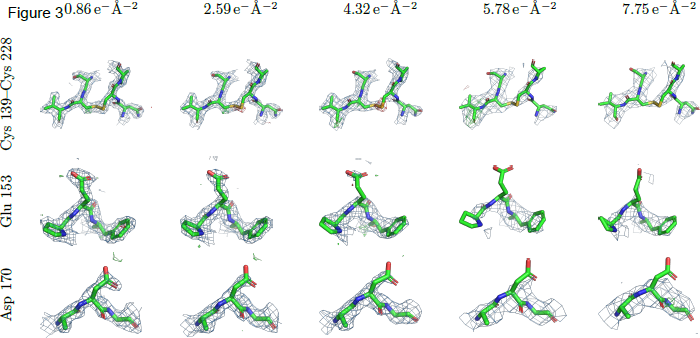
Disulfide bond breakage and decarboxylation of acidic side chains indicate site-specific radiation damage in proteinase K. 2*mF*_o_-*D*F_c_ maps (blue meshes) are contoured at 1.5 σ above the mean, *m*F_o_-*D*F_c_ difference densities (green/red meshes) are contoured at ±3 σ above/below the mean. Maps up to and including those calculated at 4.3 e^−^ Å^−2^ use data extending to 2.2 Å; the two maps at the highest exposure only use reflections up to 3.2 Å. Densities are carved to 2 Å around the selected atoms. All figures were generated using PyMol (Schrödinger LLC, 2014).

We also observe site-specific damage in the decarboxylation of the acidic side chains (Figure 3). Like the disulfide bonds, these moieties have been observed to be particularly sensitive to radiation in both X-ray crystallography (Weik et al., 2000) and single-particle cryo-EM (Bartesaghi et al., 2014; Barad et al., 2015). In proteinase K, the density around the side chains of glutamate and aspartate begins to deteriorate starting at a total exposure of ~2 e^−^ Å^−2^ and completely disappears after approximately 5 e^−^ Å^−2^ (Figure 3).

The peptide unit cell contains a zinc atom as well as an acetate and three modeled water molecules, all of which display significant signs of site-specific radiation damage at exposures >0.8 e^−^ Å^−2^ (Figure 4). For the bound zinc, the radiation damage is primarily modeled using atomic displacement parameters (ADP). Unlike occupancies, which model large-scale discrete disorder and were fixed at unity in all models, the ADP describe harmonic vibrations around the mean position of the atoms. While the density around the zinc atom in the model remains positive even at the highest exposure, its ADP steadily increases with exposure (Figure 4). The displacement begins with as little total exposure as 0.2 e^−^ Å^−2^, when the ADP of the zinc is 1.3× higher than the average ADP in the model.

**Figure 4:**
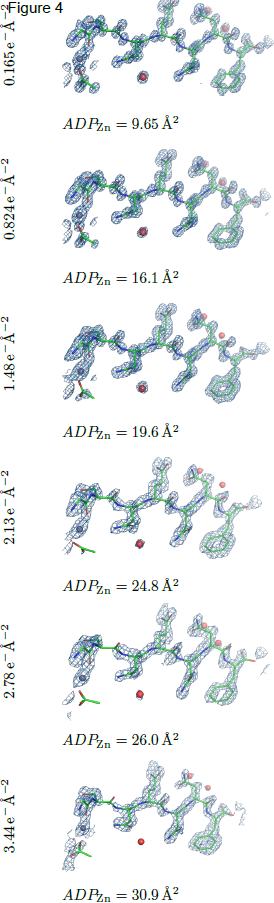
Exposure dependency on the hepta-peptide density. 2*mF*_o_-*DF*_c_ (blue meshes, countered at 1.5 σ above the mean) and *m*F_o_-*D*F_c_ density (red/green meshes, contoured at ±3σ above/below the mean) from the GSNQNNF hetpa-peptide at 0.0017 e^−^ Å^−2^ s^−1^. The maps at electron exposures up to and including 1.5 e^−^ Å^−2^ are calculated using all observed reflections to 1.01 Å; remaining maps use reflections up to 1.45 Å. The atomic displacement parameter of the Zn atom, *ADP*_Zn_, (purple sphere to the left), increases steadily over the course of exposure.

**Figure 5:**
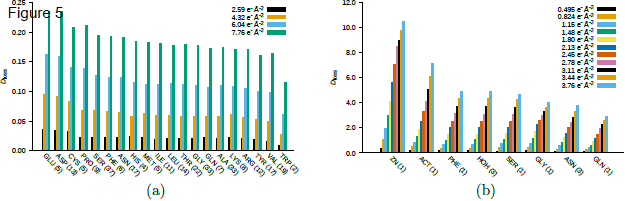
Accumulated density loss. Density loss in arbitrary units for all the amino acids, ligands, and ions present in the refined models of (a) proteinase K and (b) the hepta-peptide. The entities are sorted in the approximate order of initial damage onset. The number in parentheses denotes the occurrence of the respective amino acid in the structure. Only reflections in the resolution range common to all datasets are considered (20.8–3.20 Å for proteinase K; 14.0–1.45 Å for GSNQNNF).

Site specific damage was further assessed in real space using RIDL (Bury et al., 2015), which calculates the maximum density loss *D*_loss_ for each atom in the model. This provides a means to objectively establish the sensitivity of the different amino acids to electron radiation. Generally, the results from MicroED confirm the order and appearance of site-specific damage observed in X-ray crystallography: metals are significantly more prone to damage and glutamate, aspartate, and cysteine residues accumulate damage even at very low exposures to the electron beam.

## Discussion

Past studies of radiation damage in cryo-EM described the exposure-dependent decay of diffraction intensities up to and including 3 Å resolution (Henderson & Unwin, 1975; Hayward & Glaeser, 1979; Jeng & Chiu, 1984; Baker et al., 2010). No data was analyzed at resolutions better than 3 Å likely because such data was not recorded at the time. Unfortunately, even a recent study that reported a 2.6Å resolution single particle reconstruction included an analysis of beam induced damage only to lower resolution (Grant & Grigorieff, 2015). Moreover, while the early experiments analyzed the decay of diffraction intensities, little or no analysis of the effects of exposure on real space was described. This is likely because at low resolutions, the effects of specific damage in real space are difficult to characterize to any degree of accuracy.

We performed a deep analysis of the effects of electron beam radiation damage on biological samples at resolutions better than 1Å using MicroED (Shi et al., 2013; Nannenga et al., 2014b). Using ultra-low exposures allowed us to collect sufficient data from several crystals to allow structure determination. Each crystal was then sequentially exposed to the electron beam and additional structures were determined, from the very same crystals, at increasing levels of total exposure. In this way, we could investigate the effects of exposure on both reciprocal and real space informing us on both global and site-specific damage to the sample. With this approach, we could follow the trends of beam induced damage in biological matter at very high resolutions (better than 1 Å) in a way not previously possible.

Our real-space analysis shows that site-specific damage is apparent at high resolution even with exposures less than 1 e^−^ Å^−2^; these exposures are well below those currently used in other cryoEM modalities, for example imaging in single particle EM. This analysis therefore holds important implications for all cryo-EM methods, particularly single particle EM as recently resolutions that approach those commonly observed in crystallography have been reported (Merk et al., 2016).

Imaging in single particle cryo-EM allows the determination of protein structures from a collection of thousands of projection images of individual particles oriented randomly in vitrified ice. When a sufficiently fast camera is available, single-particle cryo-EM data is often collected as a movie. The exposure delivered to the sample reflects a tradeoff between contrast and loss of high-resolution information to radiation damage. Using catalase crystals, it was previously suggested that the optimal tradeoff between signal and damage was ~20 e^−^ Å^−2^ for the target resolution of 20Å, while for 3Å resolution it was recommended that 10 e^−^ Å^−2^ be used (Baker et al., 2010). The total exposure, typically more than 20 e^−^ Å-^2^, is fractionated over a sequence of short exposures. This allows individual frames to be corrected for specimen drift and beam-induced movement prior to averaging, while the first frame is usually discarded during processing. The last frames aid alignment, but are excluded from the final average because they contribute little high-resolution information. The average of the first 2 to ~5 frames from the movie therefore reflect a superposition of the same particles exposed to electron doses in the range of 2 e^−^ Å^−2^ to 10 e^−^ Å^−2^ and these are used for the final reconstruction when combined with data from thousands of other particles.

Since the damage mechanisms in all of these EM methods originate from the same phenomena, it is likely that the effects of randomly distributed damage events are washed out during the immense averaging and the use of methods to exclude certain particles from the final reconstruction in single particle EM. However, site-specific damage to acidic side chains has already been observed at 3.2 Å in 10 e^−^ Å^−2^ exposures (Bartesaghi et al., 2014).We surmise that the underlying damage to the sample will limit the attainable resolution in single-particle cryo-EM at the doses currently used (Grant & Grigorieff, 2015; Merk et al., 2016) and that with such exposure levels the collected data is of mostly damaged particles. When known structures are reconstructed by single particle EM, even if the resulting map is noisy because of damage, it can still be readily interpreted because the correct answer is available. But for novel structures, where the correct structure is unknown, building structures de novo with noisy maps of damaged protein is very challenging and could be prohibitive.

Quantification of radiation damage and estimates of crystal lifetime under irradiation not only depend on the sample (e.g. the number of scattering unit cells and their size, composition and thickness of the surrounding mother liquor and the embedding vitrified ice) and the measurement setup (e.g. quanta, dose, and temperature) but also on how data were processed and analyzed. Measures such as the upper resolution limit, *B* factors, and unit cell volume are often the result of some optimization procedure and may be affected by factors other than the actual damage to the crystal (Kmetko et al., 2006). This is reflected in the literature by the wide spread of dose limits. The *D*_50_ value of 2.2 e^−^ Å^−2^ calculated from reflections of proteinase K in the 21.0–1.7 Å interval is consistent with past measurements using electron diffraction from two-dimensional crystals (Stark et al., 1996) as well as three-dimensional crystals (Unwin & Henderson, 1975; Jeng & Chiu, 1984; Baker et al., 2010). When *D*_50_ was calculated from the reflections in the 14.0–1.7 Å interval of the much smaller hepta-petide, its value was 2.0 e^−^ Å^−2^ and 2.2 e^−^ Å^−2^ for high and low exposures, respectively, very close to the value obtained from proteinase K. In line with previous studies in synchrotron X-ray crystallography at comparable flux densities and temperatures, we do not see any effects from the dose rate on the observed global damage (Holton, 2009; Warkentin et al., 2013).

Given that the absorbed dose will ultimately limit the amount of meaningful data that can be extracted from a sample, data collection in the face of radiation damage may be viewed as an optimization problem. The more electrons are delivered to the sample in cryo-EM the stronger the signal, but then noise and damage accumulate and the high-resolution information suffers. Where each sample need only be exposed once, e.g. single-particle cryo-EM or serial femtosecond crystallography (Schlichting, 2015), the exposure can be tuned to maximize the signal-to-noise ratio. When pictures are recorded in cryo-EM a minimum level of exposure is required so that sufficient signal is recorded on the camera to facilitate phase contrast and faithful reconstructions. This need for phase contrast makes it hard to lower the total dose in these experiments where it was recommended to use 10 e^−^ Å^−2^ for optimal tradeoff between signal and noise for the target resolution of only 3 Å (Baker et al., 2010).

In MicroED, where phases are lost and only amplitudes are recorded, the minimum exposure necessary for recording the signal is significantly lower than in single particle EM. This allows data collection at extremely low exposures <0.01 e^−^ Å^−2^ s^−1^ and entire data sets to be collected from a total exposure less than a single electron per Å^2^ at which point atomic resolution information can be preserved. Our estimated *D*_50_ rate of approximately 2 e^−^ Å^−2^ sets an upper target for single particle cryo-EM experiments seeking to maximize resolution. This is particularly challenging but with increasingly sensitive cameras and better algorithms to allow the use of the first few frames of the recorded movies one might be able to achieve such a feat.

## Materials and methods

Frozen-hydrated crystals of proteinase K and a 7-residue peptide were prepared as previously described (Shi et al., 2016; Martynowycz et al., 2017), except the blotting force and time of the FEI (now Thermal Fischer) Vitrobot Mark IV were optimized to position 24 and 12 s, respectively, at an environment humidity of 30% for proteinase K and 20 and 20 s for GSNQNNF. Electron diffraction datasets from separate crystals were collected using an FEI Tecnai F20 transmission electron microscope operated at 200 kV, with the objective aperture fully open to evenly illuminate an area extending beyond the sample and setting the selected area aperture to closely match the size of the crystal. A frozen-hydrated grid was loaded onto a Gatan 626 cryo-holder and transferred to the microscope, where the specimen temperature was maintained at ~100 K. All diffraction images were acquired using a TVIPS TemCam-F416 CMOS camera and corrected to account for negative pixel values (Hattne et al., 2016) prior to further processing.

### Proteinase K data collection and processing

For each crystal of thickness 200–400 nm, the same 23° wedge was repeatedly collected up to five times by continuously rotating the stage from −12° to +11° (−38° to −15° for crystal 3) off its untilted orientation at a constant rate of 0.089° s^−1^ (Nannenga et al., 2014b). The rate of electron exposure was adjusted to 0.007 e^−^ Å^−2^ s^−1^., calibrated using a Faraday cage. The individual datasets, each consisting of 49–50 frames with exposure time 5.1 s were recorded at a camera length of 1.2 m, corresponding to a virtual detector distance of 2.2 m.

Proteinase K data were indexed and integrated in *P*4_3_2_1_2 using MOSFLM (Leslie & Powell, 2007). Wedges from six different crystals were merged by the order in which they were collected using AIMLESS (Evans & Murshudov, 2013), and the set of free reflections was copied from the molecular replacement search model, PDB entry 5i9s. Neither of these crystals yield a complete dataset on their own, but since proteinase K does not exhibit any pronounced preferred orientation, this produced five reasonably complete datasets, each comprised of frames with a similar degree of exposure. The choice between intensities derived from summation integration and profile fitting was left up to the optimization algorithm implemented in AIMLESS; in all cases this resulted in profile-fitted intensities being used. Relative *B*-factors were calculated between merged single-crystal datasets with SCALEIT from the CCP4 suite (Howell & Smith, 1992; Winn et al., 2011).

To quantify these global effects of radiation damage, integrated intensities were averaged for each diffraction image. Averages from different crystals were scaled by a single factor in the range [0.10, 1.0] and simultaneously fit to a common function on the form *A* × exp(−*B*×*x*) using the BFGS minimizer implemented in scipy (Oliphant, 2007). The mean effective resolution was calculated by EFRESOL (Urzhumtseva et al., 2013) and used as an objective high-resolution cutoff for all datasets.

The first dataset was phased by molecular replacement using MOLREP (Vagin & Teplyakov, 1997) from PDB entry 5i9s, and the solution was reused for all subsequent datasets. Water molecules and ions were excluded from the refined structure: while these improve the quality of the model at high resolution, they are difficult to reliably model once damage degraded the quality of the data. This model was also used to calculate solvent-accessible areas with AREAIMOL (Winn et al., 2011).

All models were refined with REFMAC (Murshudov et al., 2011), with electron scattering factors calculated using the Mott–Bethe formula. The occupancies were set to unity for all atoms and no alternate confirmations were used to model partial damage to specific sites of the molecule. Further processing and refinement statistics are given in Table 1.

### GSNQNNF data collection and processing

Crystals of the hepta-peptide GSNQNNF that were 100–500 nm thick, were tilted over ~60° at a three-fold higher rotation rate (0.3°s^−1^) than was used for proteinase K and up to 12 sweeps were collected from each crystal. To probe the effect of dose rate on radiation damage, peptide data were collected at 0.0028 e^−^ Å^−2^ s^−1^ and 0.0017 e^−^ Å^−2^ s^−1^. These rates were tuned to maximize the number of sweeps collected from an individual crystal. Single crystal datasets comprised of approximately 100 images were collected with an exposure time of 2.1 s and camera length 0.73 m which corresponds to an effective sample to detector distance of 1.2 m. Because two orders of magnitude fewer reflections are typically observed on a diffraction pattern from short segments like GSNQNNF than from proteinase K, intensities were averaged for each dataset instead of for each frame when estimating the effects of global damage on the hepta-peptide.

The datasets were indexed and integrated in *P*1 with XDS (Kabsch, 2010b) and an isomorphous subset of 10 and 12 crystals for the peptide at 0.0028 e^−^ Å^−2^ s^−1^ and 0.0017 e^−^ Å^−2^ s^−1^, respectively, was scaled and merged with XSCALE (Kabsch, 2010a). Phases for the GSNQNNF data were determined *ab initio* by direct methods from the first collected data set using SHELXD (Sheldrick, 2008). XDSCONV (Kabsch, 2010b) was used on this dataset to assign a free set of reflections, which was subsequently reused for all later peptide datasets. A ligated acetate, three water molecules, and a single zinc atom were included with the GSNQNNF model, because they constitute a significant fraction of the unit cell contents, and all atoms were fixed at full occupancy. Otherwise data collection and processing were performed as detailed for proteinase K; statistics for the datasets at the high and low dose rates are given in Table 2 and Table 3, respectively.

## Acknowledgements

The Gonen laboratory is supported by funds from the Howard Hughes Medical Institute. J.A.R. is partly supported by NSF Grant No. DMR 1548924, DOE Grant DE-FC02-02ER63421, and as a Searle Scholar and a Beckman Young Investigator.

## Author Contributions

DS, JAR, and TG designed the experiment; DS, CG, and CZ collected data; JH, CZ, MG-J, and MWM analyzed data; JH, DS, JAR, and TG wrote the manuscript with input from all authors.

